# SoxS represses flagellar gene expression through RflP in *Salmonella enterica* Serovar Typhimurium

**DOI:** 10.1101/2020.05.29.124750

**Authors:** Srinivas S. Thota, Brittany N. Henry, Lon M. Chubiz

## Abstract

Flagellar gene expression is subject to regulation by many global transcription factors in response to environmental and nutritional signals. One of the primary ways this occurs in *Salmonella enterica* serovar Typhimurium, and its close relatives, is through controlling levels of FlhD_4_C_2_ (the flagellar master regulator) via transcriptional, post-transcriptional, and post-translational mechanisms. Recently, we found the homologous transcription factors MarA, Rob, and SoxS repress *flhDC* expression by distinct mechanisms. MarA and Rob, regulators involved in inducible multidrug resistance, repressed *flhDC* transcription by interacting directly with the *flhDC* promoter. Alternatively, SoxS, the oxidative stress response regulator, altered FlhD_4_C_2_ levels independent of *flhDC* transcription by post-transcriptional or post-translational mechanism. Here, using a forward genetic screen, we discovered that SoxS-dependent repression of flagellar gene expression occurs through RflP, an anti-FlhD_4_C_2_ factor that targets FlhD_4_C_2_ for proteolytic degradation. Elevated *soxS* expression resulted in concomitant increases in *rflP* expression, indicating SoxS may work through RflP at the level of *rflP* transcription. Mapping of the *rflP* promoter and a bioinformatic search yielded a putative SoxS binding site proximal to the *rflP* transcription start site. Comparison of the *rflP* promoter region in *S*. Typhimurium and *Escherichia coli* indicate substantial differences, providing a possible mechanism for differential expression of *rflP* between these species.

**IMPORTANCE:** *Salmonella enterica* is a major cause of foodborne illness. Understanding environmental and intracellular signals used by *Salmonella* to control expression of virulence-associated traits is critical to advancing treatment and prevention of *Salmonella*-related disease. Reduced expression of flagella at key points during *Salmonella* infection aids in evasion of the host innate immune system. Within macrophages *Salmonella* is non-flagellated and exposed to oxidative stress. SoxS-dependent repression of flagellar genes may provide a potential link between oxidative stress and reductions in flagellar expression.

## INTRODUCTION

Flagellar motility is a primary means of locomotion for many bacterial species. While this form of motility allows migration to more favorable environments, the production and operation of flagella are significant biosynthetic and energetic burdens for flagellated bacteria (1, 2). For this reason, most bacteria tightly regulate the biosynthesis of flagella in response to diverse environmental cues (3, 4).

In *Salmonella enterica* serovar Typhimurium, and related *Enterobacteriaceae*, flagellar genes form a distinct regulon where expression is hierarchical (3, 5). In this arrangement, flagellar genes are temporally expressed in three distinct classes that correlate to specific stages of flagellar assembly. Atop the hierarchical flagellar regulon is the heterohexameric transcription factor FlhD_4_C_2_, the product of the Class I operon *flhDC (6*). FlhD_4_C_2_ is considered the master regulator that activates expression of several Class II genes corresponding to components of the flagellar basal body and includes an alternative sigma factor, FliA (or σ^28^) (7–9). FliA subsequently directs transcription of an array of Class III genes encoding the flagellar motor and filament, as well as proteins involved in chemotaxis (10). Additionally, a number of internal regulatory feedbacks exist to ensure sequential expression of Class II and Class III genes including autorepression of *flhDC* expression by FlhD_4_C_2_, FliT-dependent inhibition of FlhD_4_C_2_ activity, indirect activation of FlhD_4_C_2_ activity by FliZ, and FlgM (a secretion-dependent anti-FliA factor) inactivation of FliA (11–18).

Apart from autogenous control, numerous environmental and nutritional signals regulate flagellar gene expression. Integration of these signals, and the resulting regulation of flagellar gene expression, occurs primarily through control of *flhDC* expression or by altering the stability of FlhD_4_C_2_. At the level of *flhDC* transcription, several global transcription factors involved in nutrient and envelope stress sensing serve as activators and repressors including: CRP, Fis, Fur, H-NS, RcsB, and OmpR; among several others (11, 19–24). Additionally, the homologous multidrug resistance-associated transcription factors MarA and Rob were recently found to directly repress *flhDC* transcription in *S*. Typhimurium (25). Beyond transcription-level control, a number of small regulatory RNAs (sRNA) are known to control *flhDC* expression at the mRNA level (26). Together, these mechanisms carefully titrate the concencentration of FlhD_4_C_2_ produced across variable environments and growth stages.

While transcriptional and post-transcriptional mechanisms control FlhD_4_C_2_ production, protein-protein interactions between RflP (also known as YdiV) and FlhD_4_C_2_ are known to regulate FlhD_4_C_2_ levels by targeting FlhD_4_C_2_ for degradation by ClpXP protease (27). The effects of RflP on FlhD_4_C_2_ levels are strongly nutrient dependent (28–30). Despite being an EAL-domain containing protein, RflP has not been demonstrated to have dicyclic-GMP phosphodiesterase activity (28, 31, 32). Thus, RflP regulation of FlhD_4_C_2_ abundance occurs through changes in RflP concentrations dictated by *rflP* transcription (15, 28–30, 33). Detailed mapping of *rflP* regulation remains cryptic (15). However, nutrient and envelope stress have been observed to increase *rflP* expression (28, 34). Interestingly, despite being conserved in flagellated *Enterobacteriaceae*, the effects of RflP appear to be variable across and within species (29). For example, RflP strongly regulates flagellar expression in *S*. Typhimurium and pathogenic *E. coli* strains, but not in the laboratory model *E. coli* K-12 (29, 35). The variability in RflP usage is likely a result of differences in *rflP* regulation, but precise mechanisms remain largely unexplored.

The superoxide stress response regulator SoxS is a potent repressor of flagellar gene expression (25). Similar to its homologs MarA and Rob, SoxS is an AraC-family transcription factor (36). Levels of SoxS are controlled through *soxS* transcriptional activation by the redox-sensing transcription factor SoxR and proteolytic degradation of SoxS by Lon protease (37–39). SoxS is responsible for directly and indirectly activating and repressing over 30 genes in *E. coli* (40, 41). Many of the genes regulated by SoxS code for proteins involved in detoxifying intracellular superoxide, efflux of toxic compounds, decreasing membrane permeability, and altering central carbon metabolism (40, 42). More recently, SoxS was observed to repress expression of the flagellar regulon *S*. Typhimurium (25). SoxS-dependent flagellar repression occurred through reductions in *flhDC* transcription and post-transcriptional repression of FlhD_4_C_2_ production in an Hfq-independent manner. The identity of factors involved in SoxS-dependent repression of flagellar gene expression are unknown.

Here, we have discovered SoxS represses flagellar gene expression by reducing FlhD_4_C_2_ concentrations via increased expression of *rflP*. Using a transposon-based forward genetic screen, we identified several independent insertion mutants in the *rflP* coding and promoter regions. Disruption of *rflP* obviated SoxS-dependent repression of Class II and Class III gene expression, restoring motility. The regulatory connection between SoxS and *rflP* appears to be unique in *Salmonella*. The ability of SoxS to increase *rflP* expression and reduce flagellar gene expression may have a broader role in the virulence life cycle of *Salmonella* where flagellation is regulated in response to conditions within the host, particularly the presence of strong oxidizing agents.

## RESULTS

### SoxS-dependent repression of *flhDC* expression occurs primarily through a post-transcriptional mechanism

We have previously shown SoxS represses all three classes of flagellar genes (25). Similar effects on flagellar gene expression were observed for the SoxS homologs MarA, Rob, and RamA. Specifically, when SoxS was overproduced we found it repressed *flhDC* expression at the transcriptional and post-transcriptional levels (25). To better understand the effects of endogenous levels of SoxS, here, we used a constitutively active *soxR* mutant (*soxR*^Con^) that expresses *soxS* at roughly 50% of its maximally induced levels (25, 43, 44). A constitutive mutant was utilized to avoid any possible pleiotropic effects of SoxR activating compounds on flagellar gene expression. Using the *soxR*^Con^ mutant, we measured the effects of SoxS on flagellar gene transcription, levels of FlhC and FliC proteins, and motility (**Figure 1**).

**Figure 1.**
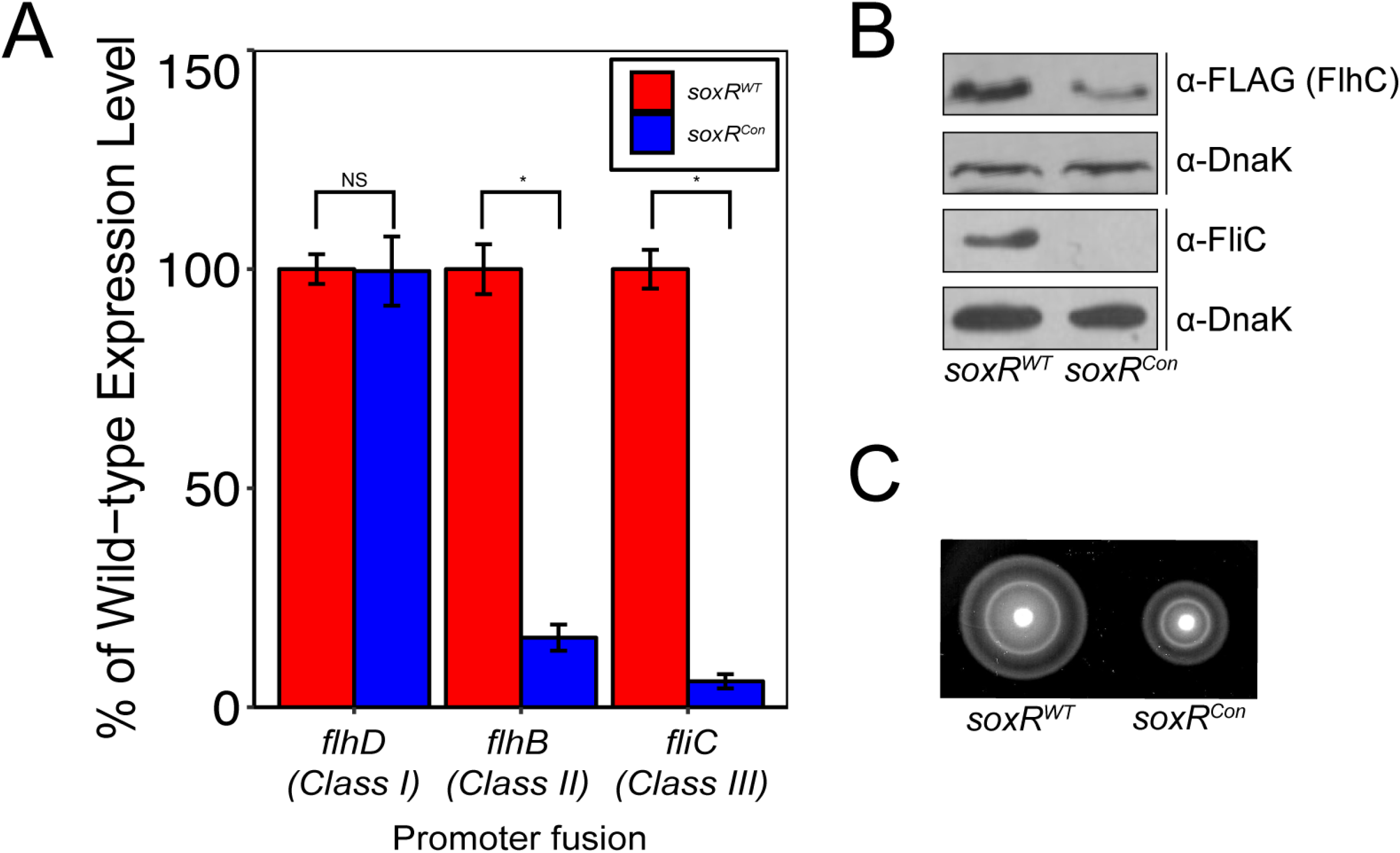
SoxS-dependent repression of flagellar genes and motility occurs after Class I transcription. **A**) Levels of of *flhDC* (LCM2324, LCM2471), *flhB* (LCM2325, LCM2472) and *fliC* (LCM2326, LCM2473) transcription in *soxR*^Con^ and WT background strains, respectively. Data are presented as the percentage of the wild-type expression level for each class of flagellar promoter. (*) indicates Student’s t-test P≤10^−6^. **B)** The effect of constitutive expression of SoxS (*soxR*^Con^) on FlhC and FliC protein levels. Western blots were conducted with 100 μg total protein. Anti-FLAG antibody was used to detect FlhC-3XFLAG expressed in *soxR*^Con^ (LCM2697) and *soxR*^WT^ (LCM2696) background strains, respectively. Anti-FliC antibody was used to detect the flagellin protein, FliC in *soxR*^Con^ (LCM2449) and *soxR*^WT^ (LCM1930) background strains, respectively. The loading control DnaK was detected by using anti-DnaK antibody. **C**) Swimming motility in *soxR*^WT^ (LCM1930) and *soxR*^Con^ (LCM2449) background strains, respectively.

We measured transcription of *flhDC*, *flhB* and *fliC* as representatives of Class I, Class II and Class III flagellar genes, respectively, in *soxR*^Con^ and *soxR*^WT^ backgrounds (**Figure 1A**). Transcription from each promoter was measured using chromosomally-integrated promoter fusions to *yfp*(Venus) similar to Koirala and coworkers (45). We found transcription of *flhB* and *fliC* were reduced to 15.9±2.9% (Student’s *t*-test, *P*=1.1×10^−8^) and 5.9±1.6% (Student’s *t*-test, *P*=1.6×10^−7^) in the *soxR*^Con^ genetic background compared to *soxR*^WT^. Interestingly, there was no significant change in the transcription of *flhDC* under the same conditions. These results are consistent with SoxS activating a post-transcriptional mechanism to repress *flhDC* expression.

Reductions in Class II and Class III transcription were reflected in decreased levels of FliC protein and motility in the *soxR*^Con^ background compared to *soxR*^WT^. Levels of FliC were qualitatively reduced below detectable limits in the *soxR*^Con^ background (**Figure 1B**). We also observed qualitative reductions in levels of FlhC protein in the *soxR*^Con^ background compared to *soxR*^WT^, providing a likely reason for reductions in FliC (**Figure 1B**). Similarly, swimming motility of the *soxR*^Con^ mutant was reduced to 66.3±5.0% of wild-type (**Figure 1C**; Student’s *t*-test, *P*=3.5×10^−12^). Taken together, the absence of changes in *flhDC* transcription and decreased levels of FlhC protein levels in response to endogenous levels of SoxS suggest SoxS represses *flhDC* expression at the post-transcriptional or post-translational levels.

### Identification of mutants that bypass SoxS-dependent flagellar repression

We employed a transposon-based forward genetic screen in the *soxR*^Con^ background for mutants with recovered flagellar gene expression. Previously, we demonstrated *in vivo* interactions between SoxS and the *flhDC* promoter are weak and may have limited biological significance (25). Here, we showed constitutive expression of SoxS at endogenous levels represses *flhDC* post-transcriptionally (**Figure 1**). These lines of evidence point to SoxS regulating *flhDC* expression indirectly through another pathway. To identify pathways involved in SoxS-dependent flagellar repression we performed a forward genetic screen in our *soxR*^Con^ background using the T-POP transposon, similar to the approach used by Wozniak and coworkers to identify flagellar regulators (46). In total, >100,000 mutants from 6 independent pools were screened for restoration of *fliC* transcription. The unique mutants recovered with restored *fliC* expression were 6 insertions in *rflP* (an anti-FlhD_4_C_2_ factor) (**Figure 2A**), 3 insertions in *fliD* (the flagellar filament cap protein), 2 insertions in *rcsB* (the Rcs response regulator); and 1 insertion each in *topA* (topoisomerase I), *prgH* (a SPI1 type III secretion structural protein), and *fliC* (the flagellar filament protein), respectively. Remarkably, we observed no T-POP insertions in the *soxRS* locus. However, the extensive T-POP screen by Wozniak and coworkers did not uncover *soxS* as a flagellar regulator, as well; suggesting the *soxRS* locus may be a cold spot for the T-POP transposon system (46).

**Figure 2.**
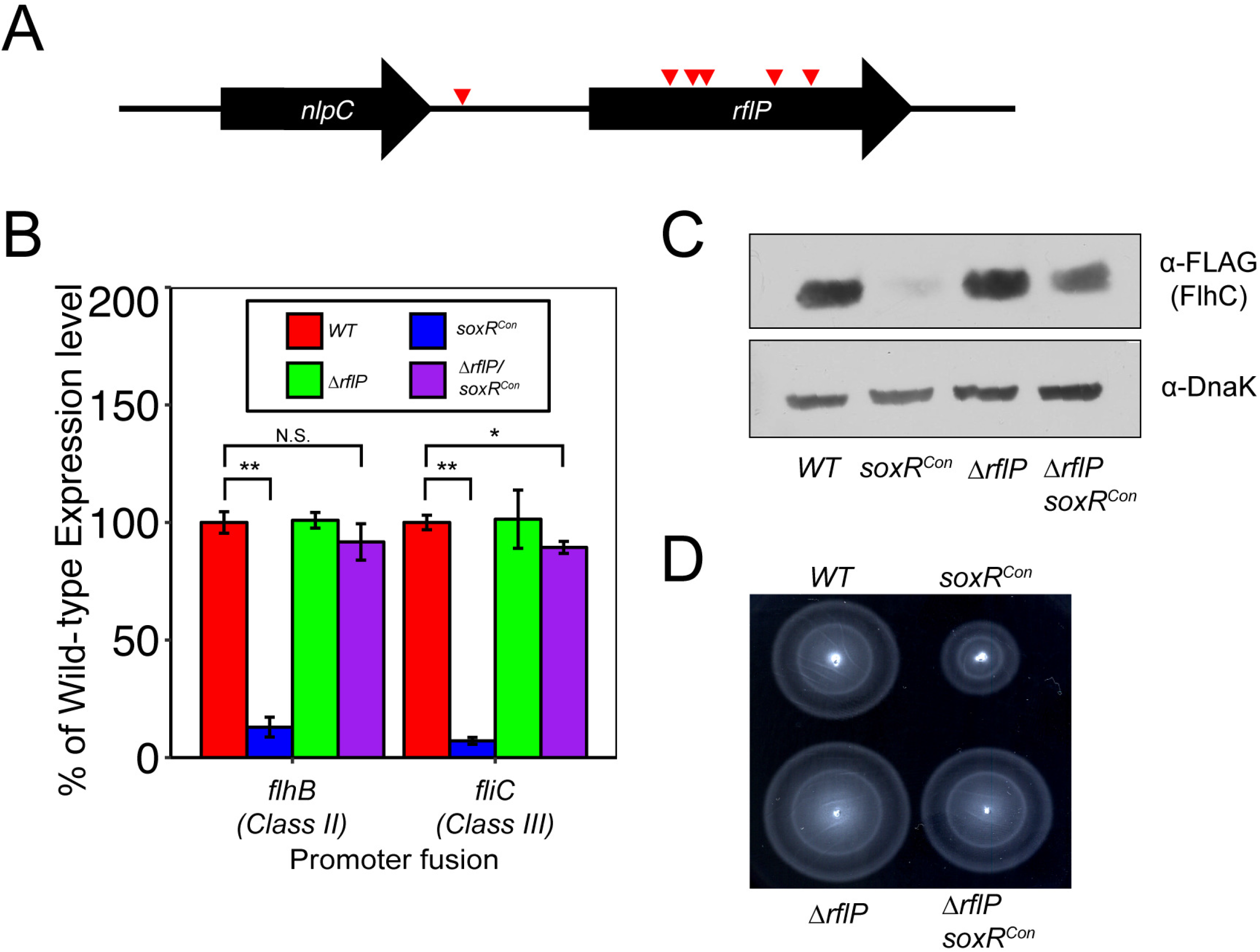
Inactivation of *rflP* alleviates SoxS-dependent flagellar repression. **A)** The relative location of T-POP insertion sites in the *rflP* gene. Insertions are indicated by red arrows. T-POP insertions in *rflP* were identified at genome coordinates 1,422,579; 1,423,247; 1,423,428; 1,423,238; 1,423,248; and 1,423,344bp (ordered from left to right). **B)** Expression of *flhB* (Class II) and *fliC* (Class III) genes in WT (LCM2325, LCM2326), *soxR*^Con^ (LCM2472, LCM2473), *ΔrflP* (LCM2808, LCM2761) and *soxR*^Con^*/ΔrflP* (LCM2809, LCM2762) background strains, respectively. Data are presented as the percentage of the wild-type expression level for each class of flagellar promoter. (*, **) indicates a Student’s *t*-test *P≤0.002* or *P*≤10^−6^. **C)** FlhC-3XFLAG protein levels in WT (LCM2764), *soxR*^Con^ (LCM2765), *ΔrflP* (LCM2766) and *soxR*^Con^*/ΔrflP* (LCM2767) background strains. Anti-FLAG antibody was used to detect FlhC-3XFLAG and anti-DnaK antibody was used to detect the loading control, DnaK. **D)** Swimming motility observed in WT (LCM1930), *soxR*^Con^ (LCM2449), *ΔrflP* (LCM2760), *soxR*^Con^*/ΔrflP* (LCM2763) background strains.

The recovered T-POP insertion mutations fell into two major categories. First were flagellar genes with possible autogenous regulatory effects on flagellar gene expression. For example, insertions in *fliD* likely have polar effects on *fliT*, a gene distal to *fliD* in the same Class II operon (*fliDST*). FliT is a chaperone for several flagellar structural proteins and also serves as a negative regulator of FlhD_4_C_2_ activity via protein-protein interactions (13). Loss of *fliT* is known to increase expression of Class II and Class III genes (12, 13). Thus, removal of a FlhD_4_C_2_ inhibitor may increase expression of *fliC*. Since SoxS is likely utilizing a pathway external to the flagellar regulon, we did not prioritize further analysis of *fliD* insertions. Second were genes with autonomous roles in flagellar gene regulation. Specifically, *rflP* and *rcsB* have known interactions with FlhD_4_C_2_ and the *flhDC* promoter, respectively. These negative interactions reduce Class II activation (23, 27, 28, 47). From the second category, we looked to examine the effects of *rflP* as its effects on FlhD_4_C_2_ are post-translational, consistent with our observations surrounding SoxS-dependent repression of Class I expression (27).

### Inactivation of *rflP* alleviates SoxS-dependent repression of flagellar gene expression and motility

We tested whether SoxS repressed flagellar genes through RflP by observing the effects of a *rflP* mutant on flagellar gene expression, levels of FlhC protein, and motility in the *soxR*^Con^ genetic background. Similar to previous observations, the expression of *flhB* (Class II) and *fliC* (Class III) flagellar genes were reduced to 12.9±4.2% (Student’s *t*-test, *P*=1.5×10^−9^) and 7.0±1.4% (Student’s *t*-test, *P*=9.6×10^−9^) of the wild-type levels, respectively, in the *soxR*^Con^ strain (Figure 2B). Inactivation of *rflP* resulted in rescue of *flhB* (91.7±7.7%, Student’s *t*-test, P=0.14) and *fliC* (89.3±2.5%, Student’s *t*-test, P=2.0×10^−3^) expression to nearly wild-type levels in the *rflP soxR*^Con^ background strain (**Figure 2B**). Likewise, the levels of FlhC protein were reduced in *soxR*^Con^, as expected, but were qualitatively recovered in the *rflP soxR*^Con^ genetic background (**Figure 2C**). Notably, the levels of FlhC in the *rflP soxR*^Con^ background, while increased compared to *soxR*^Con^, remain qualitatively lower than wild-type FlhC levels. This might be due to the presence of an additional SoxS-dependent pathway affecting FlhD_4_C_2_ stability, or the elevated levels of SoxS expression may indirectly increase FlhD_4_C_2_ degradation rates. Consistent with recovery of Class II/III gene expression and FlhC levels, the 38.4±3.7% (Student’s *t*-test, *P*=2.2×10^−6^) reduction in swimming motility in the *soxR*^Con^ background was recovered to near wild-type levels in the *rflP soxR*^Con^ background (**Figure 2D**).

We also wanted to determine if SoxS-dependent repression of flagellar genes can be complemented in the *rflP soxR*^Con^ strain when *rflP* is provided *in trans*. The *rflP* locus, including the native *rflP* promoter (described below), was introduced into the *soxR*^Con^ and *rflP soxR*^Con^ background on a low-copy number plasmid (pRflP). When harboring pRflP, the *soxR*^Con^ and *rflP soxR*^Con^ backgrounds expressed *fliC* at 8.5±0.6% (Student’s *t*-test, *P*=6.1×10^−6^) and 13.3±0.9% (Student’s *t*-test, *P*=9.4×10^−6^) of levels observed in the *soxR*^Con^ strain harboring the empty vector (pWKS30) (**Figure 3A**). Consistent with SoxS working through *rflP*, the level of *fliC* expression in the *rflP soxR*^Con^ background is 15.2±2.2 fold (Student’s *t*-test, *P*=2.4×10^−6^) more than that of *soxR*^Con^ strain bearing the control vector. Motility was also repressed in *soxR*^Con^ and *rflP soxR*^Con^ background carrying pRflP compared to same strains harboring the empty vector (**Figure 3B**). Overall, these data suggest SoxS works through RflP to decrease levels of FlhD_4_C_2_. The reduction in FlhD_4_C_2_ levels is a likely explanation for reduced expression of Class II and Class III genes, ultimately manifesting in reduced motility.

**Figure 3.**
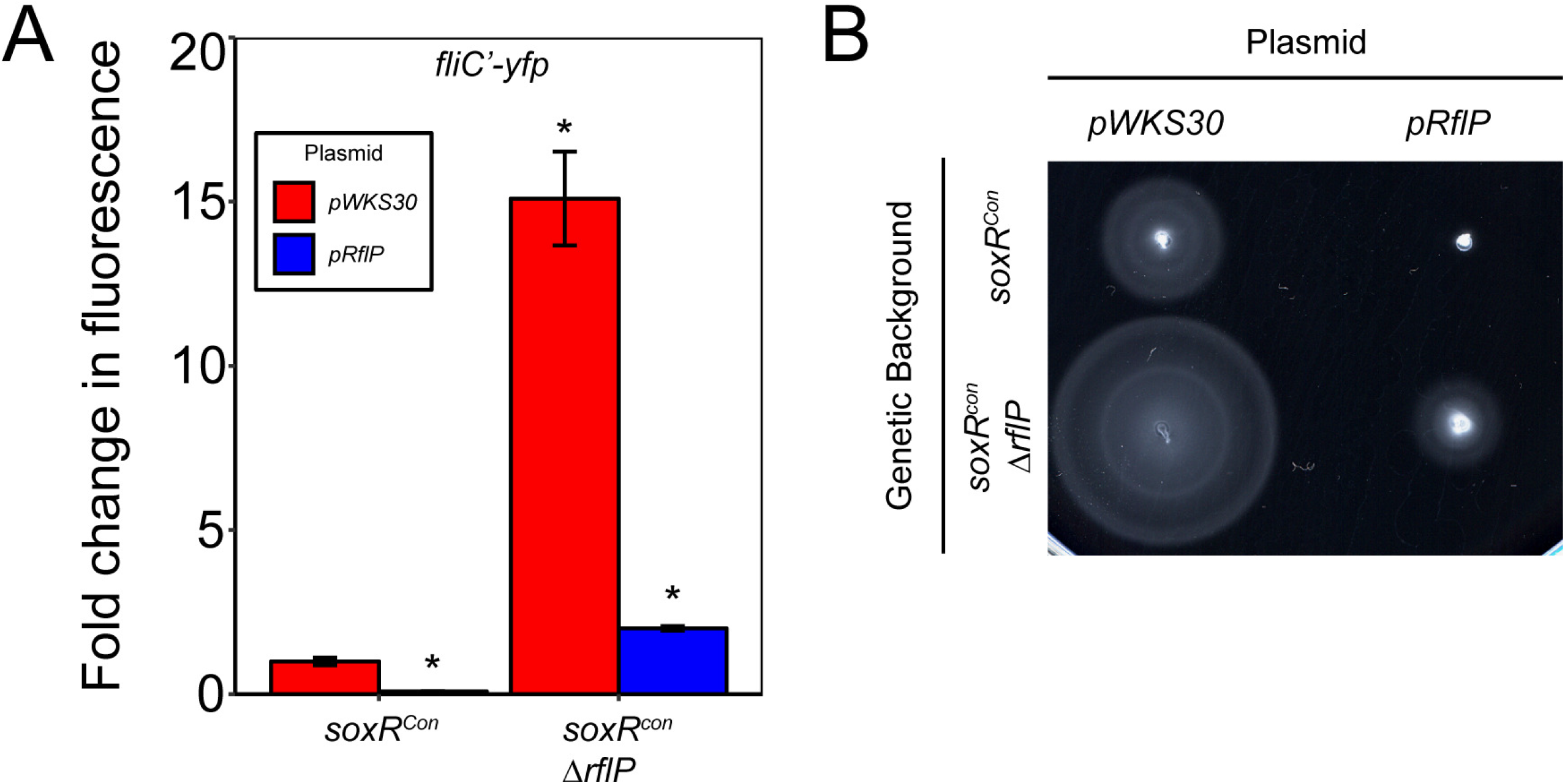
Complementation of SoxS-dependent flagellar gene repression in *soxR^Con^/ΔrflP* strain. **A)** Levels of *fliC* expression in *soxR^Con^* (LCM2473) and *soxR^Con^/ΔrflP* (LCM2762) background strains with pWKS30 (empty vector) or pRflP (*rflP* complementation vector). Data are presented as the percentage of the *fliC* expression level in the *soxR*^Con^ background harboring pWKS30. (*) indicates a Student’s *t*-test *P*≤10^−5^. B) Swimming motility in soxR^Con^ (LCM2449) and soxRCon/ΔrflP (LCM2763) strains with vectors pWKS30 and pRflP.

### Increased *soxS* expression results in elevated *rflP* promoter activity

We looked to see what effects increased *soxS* expression had on *rflP* promoter activity. One possible mechanism for SoxS to reduce flagellar gene expression through RflP is via increasing *rflP* transcription. Notably, the dosage of *rflP* expression is believed to control RflP-dependent reductions in levels of FlhD_4_C_2_ and not activation by small molecule signals, as the EAL domain of RflP is non-functional (28, 32, 33). Based on known properties of RflP-FlhD_4_C_2_ interactions and genetic evidence that SoxS functions through *rflP*, we hypothesized elevated levels of *soxS* expression may yield corresponding increases in *rflP* expression. To test this hypothesis, we used an *in situ lacZY* promoter fusion to *rflP* in *soxR*^Con^ and *soxR*^WT^ background strains. We found that *rflP* expression was 2.4±0.1 fold (Student’s *t*-test, *P*=4.8×10^−6^) higher in *soxR*^Con^ compared to wild-type (**Figure 4A**). One interpretation may be that SoxS is interacting with the *rflP* promoter and serving as an activator. The lack of transposon mutants in genes coding for transcription factors regulated by SoxS further supports this possibility. Notwithstanding, the paucity of information on *rflP* promoter structure limits any further speculation.

**Figure 4.**
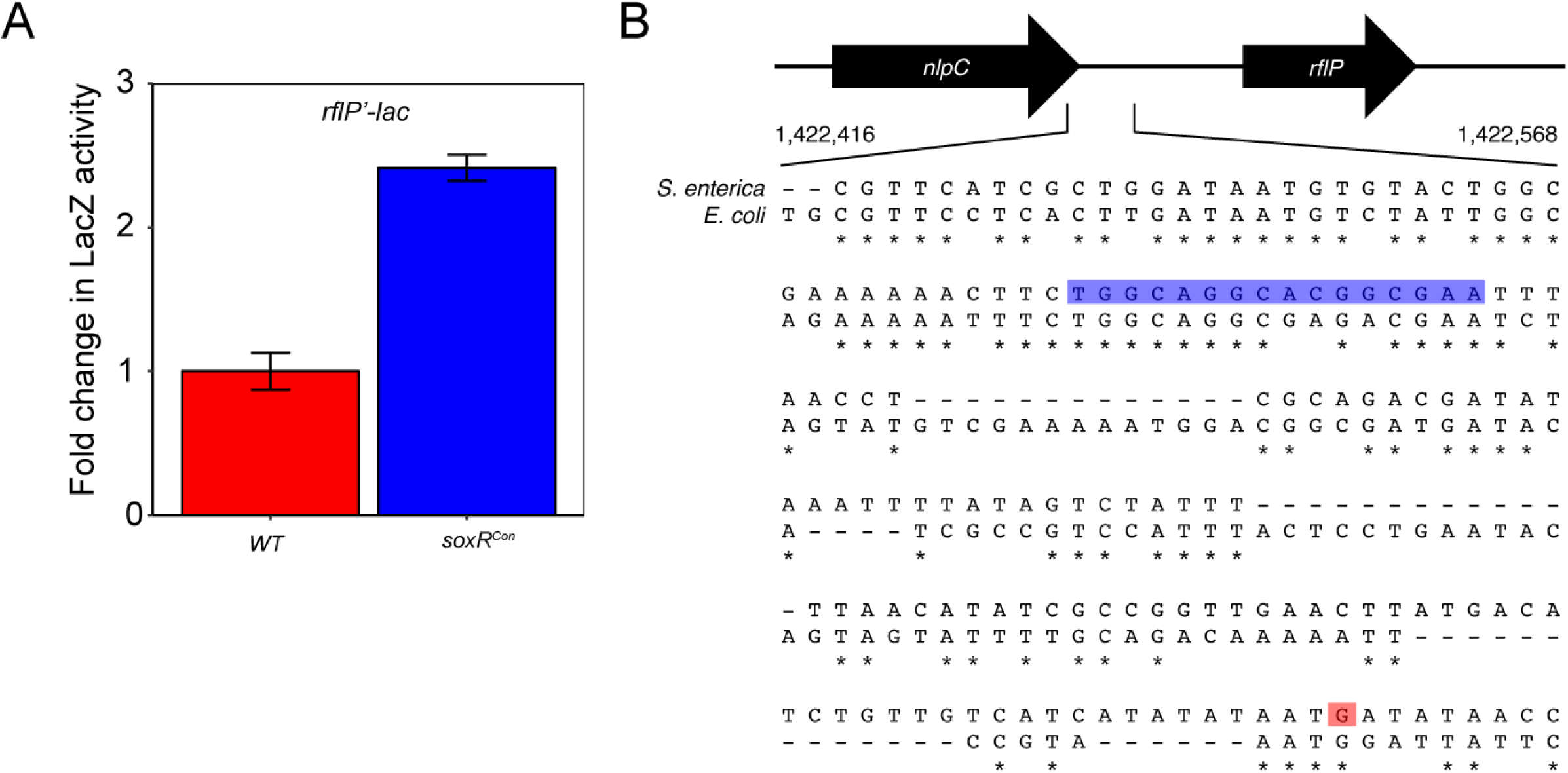
The effects of SoxS on *rflP* transcription. **A)** Levels of *rflP’-lacZY* activity in wild-type (LCM2791) and *soxR*^Con^ (LCM2792) genetic backgrounds. Data are presented as a fold increase in β-galactosidase activity compared to wild-type (LCM2791). Comparison of wild-type and *soxR*^Con^ with the Student’s *t*-test resulted in a *P*=4.8×10^−6^. **B)** The promoter region of *rflP* (1,422,425 to 1,422,575). The putative SoxS binding motif (*soxbox*) identified by MEME analysis is highlighted in blue. The guanine nucleotide highlighted in red is the transcription start site (TSS) of the *rflP* mRNA identified from 5’-RACE in wild-type.

### Mapping of the *rflP* promoter

To better understand the possible interaction between SoxS and the *rflP* promoter, we mapped the *rflP* transcriptional start site (TSS). Previous mapping of the *rflP* promoter has been via bioinformatic approaches (48, 49). Using these predictions, we found a cloned *rflP* promoter fusion cloned *in trans* displayed no response to known nutritional cues or SoxS levels (data not shown). We interpreted these data to mean that the native *rflP* promoter location may have been incorrectly mapped or may be more complex than previously predicted, as suggested by Wada and colleagues (29). Using template-switching rapid amplification of cDNA ends (5’-RACE) to sequence the 5’-end of the *rflP* mRNA, we identified the transcription start site of *rflP* to −230 bp of *rflP* start codon (**Figure 4B**). This is over 150 bp upstream of previous predictions in *E. coli* and very close to *nlpC*, the gene just proximal to *rflP* (**Figure 2A**) (48, 49). Consistent with the location of the promoter identified, in our transposon screen we only observed insertions in the *rflP* coding region or the *nlpC-rflP* intergenic region distal to the *rflP* TSS, and not in the *nlpC* gene (**Figure 2A**). This is a markedly different result than other studies in *Salmonella* that have noted disruptions in *nlpC* have polar effects on *rflP* expression (15, 29, 46). An explanation for this difference may be that SoxS utilizes the native *rflP* promoter to increase *rflP* expression while other regulators, such as FliZ, utilize the proximal *nlpC* promoter to mediate changes in *rflP* expression (15). Taken together, these data show regulation of *rflP* is complex and may involve multiple promoters.

### Identification of a putative SoxS binding site in the *rflP* promoter

Since *rflP* transcription was elevated when SoxS expression was increased, we wanted to bioinformatically search for any possible SoxS binding sites (*soxbox*) in the *rflP* promoter region. To locate possible SoxS binding sites in the *rflP* promoter, we employed the MEME motif search algorithm using 29 known and putative SoxS binding sites in *E. coli* promoters as a training data set (50). The *rflP* promoters from *S*. Typhimurium and *E. coli* were added to detect possible SoxS binding motifs. *E. coli* sequences were chosen due to the paucity of well-characterized SoxS binding sites in *S*. Typhimurium, although many of genes are similarly regulated by SoxS (51). Using these known *soxbox* containing promoters, MEME identified a conserved *soxbox* motif in the *S*. Typhimurium *rflP* (**Figure 4B**). The putative SoxS binding site (5’TGGCAGGCACGGCGAA3’) located 88 bp upstream of the *rflP* TSS is similar to *soxbox* motifs from studies by Seo and coworkers (5′-AYRGCAYWAWWTRYYAAW-3′) and Martin and workers (5’-AYNGCACNNWNNRYYAAA-3’) (41, 52). While the predicted SoxS binding site is further upstream from the TSS than *soxbox* sites in many well-studied promoters (52); examples such as the *micF* promoter indicate SoxS binding sites can be even further upstream of the TSS (*i.e*. >100 bp) (53). Future biochemical mapping of SoxS interactions with the *S*. Typhimurium *rflP* promoter will help validate these predictions.

Interestingly, this same site was predicted in the *E. coli rflP* promoter. Notably, the putative *S*. Typhimurium and *E. coli* SoxS binding sites vary in the region between the conserved 5’-GCA-3’ and 5’-YYAA-3’ minor motifs (**Figure 4B**). These differences may provide an explanation as to why SoxS has not been found to bind to the *rflP* promoter or regulate *rflP* transcription in *E. coli (41*).

### The *rflP* promoter regions of *S*. Typhimurium and *E. coli* have limited similarity

Having mapped a *rflP* promoter region by 5’-RACE in *S*. Typhimurium, we looked to compare this region to the corresponding chromosomal region in *E. coli* K-12 to determine if the *rflP* promoter was conserved. Comparison of the *rflP* promoter region in *S*. Typhimurium to the corresponding region in *E. coli* indicated substantial differences (**Figure 4B**). In particular, the region corresponding to *S*. Typhimurium *rflP* TSS bears little similarity to the same region in *E. coli* (41.3% identity between the *rflP* TSS and putative *soxbox*). This result indicates the *rflP* promoter structure may be drastically different in *E. coli* as compared to *S*. Typhimurium. In *E. coli*, genome-scale transcriptomics and chromatin immunoprecipitation sequencing (ChIP-seq) experiments involving SoxS have not identified *rflP* as a SoxS target (41). The difference in *rflP* promoter sequences may explain the absence of SoxS interactions with *rflP* in *E. coli*.

## DISCUSSION

We have demonstrated that SoxS represses flagellar gene expression primarily through elevating levels of *rflP* (also known as *ydiV*) expression. In a recent study, we established the intracellular superoxide response regulator SoxS is a repressor of flagellar genes primarily through reductions in *flhDC* (Class I) expression (25). While the homologs of SoxS, specifically MarA and Rob, were found to repress *flhDC* by direct interaction with the *flhDC* promoter, SoxS was found to decrease FlhD_4_C_2_ levels independent of effects on *flhDC* transcription (25). Post-transcriptional regulation of *flhDC* expression by SoxS was found to be independent of Hfq, the sRNA chaperone (25). This largely discounted the effects of sRNA as a possible mechanism as most sRNA that regulate *flhDC* mRNA stability and translation are known to require Hfq (54). To elucidate factors involved in SoxS-dependent post-transcriptional repression of *flhDC*, we performed a transposon-based forward genetic screen for mutants with recovered flagellar gene expression in a genetic background where SoxS is constitutively produced. Several independent insertion mutants mapped to *rflP*. Subsequent genetic analysis confirmed that *rflP* is indeed the link between increased SoxS levels and repression of flagellar gene expression. Mapping of the *rflP* transcription start site and resulting bioinformatic analysis identified a putative *soxbox* in the *Salmonella rflP* promoter region. These results shed light on a novel regulatory pathway connecting oxidative stress and flagellar gene regulation.

SoxS-dependent activation of *rflP* occurs in *S*. Typhimurium but not *E. coli* K-12 (41). Our bioinformatic analysis identified a putative *soxbox* in the *rflP* promoter in *S*. Typhimurium. The corresponding site was also present in *E. coli*; however, *E. coli* lacks the promoter region identified in *S*. Typhimurium. Transcriptomic and DNA binding studies of SoxS have indicated several common targets between *E. coli* and *Salmonella*, however, these do not include *rflP* (41, 51, 55). The strikingly different sequences of the *rflP* promoter region between these two species may provide a possible mechanism for the lack of SoxS-dependent activation of *rflP* in *E. coli* (**Figure 4B**). Further comparative studies between the SoxS regulons of these species will provide a better picture of how each species differentially uses the response to oxidative stress to alter physiology and metabolism, as demonstrated by SoxS regulation of flagellar expression.

More broadly, control of *rflP* may be an important aspect of flagellar expression in response to intracellular chemical stress in the *Enterobacteriaceae*. In species where RflP is a prominent mechanism for flagellar regulation, levels of *rflP* expression have been found to be responsive to nutritional status, envelope stress, and the cell’s physiochemical environment (28, 34, 35). Adding to the complexity of *rflP* regulation is the action of SoxS, a key mediator of the oxidative stress response in *Salmonella*. These examples highlight how *rflP* has been integrated into diverse stress responses of this bacterial family. While this study did not directly explore the interactions between SoxS and the *rflP* promoter, future studies examining the role of SoxS-dependent regulation of *rflP* throughout the *Enterobacteriaceae* will more clearly define this interaction and provide deeper insights into adaptive benefits of reducing flagellar expression in response to oxidative stress.

The relationship between SoxS and *rflP* expression could have an impact on *Salmonella* virulence. Notably, RflP-dependent reductions in flagellar expression aid in evasion of the host caspase-1 inflammatory response, enhancing the ability of *Salmonella* to survive during systemic infection (56). During this process, *Salmonella* is exposed to oxidative stress which is known to be a chemical cue for expression of certain virulence traits. The role of SoxS in this process is unclear. Earlier work by Fang and coworkers found SoxS is not involved in virulence (57). More recently, Wang and coworkers have discovered that SoxS does indeed play a regulatory role in *Salmonella* virulence (55). Specific causes for this discrepancy are unresolved but may be due to differences in the *soxS* mutants used in each study (55). While it is tempting to draw connections between SoxS, RflP, and *Salmonella*’s survival within the host, more detailed studies of the mechanism behind SoxS-dependent effects on *Salmonella* virulence are needed.

Regulation of stress responses and structural genes is complex and has diverged even amongst closely related bacterial species such as *S*. Typhimurium and *E. coli (58, 59*). In *S*. Typhimurium we have previously shown that the multidrug resistance regulators MarA, SoxS, and Rob repress flagellar gene expression in contrast to what has been observed in *E. coli* (25). Building on this, we have shown here that even amongst homologous transcription factors there exists different mechanisms to achieve the same ends. SoxS-dependent repression of flagellar expression through RflP is so far unique to *S*. Typhimurium. Further exploration of how RflP is used in flagellar regulation in other motile *Enterobacteriaceae* may shed light on how the costs and benefits of flagellar motility are managed in response to the complex environment inhabited by this bacterial family.

## MATERIALS AND METHODS

### Bacterial strains, plasmids and media

All the bacterial strains and plasmids generated in this study are listed in **Table S1**. For all experiments, cultures were propagated in tryptone broth (1% tryptone, 0.8% NaCl) and grown at 37°C Swimming motility assays were performed in soft agar (0.3% Bacto agar, 1% tryptone, and 0.8% NaCl). All genetic manipulations were conducted with cells grown in Luria-Bertani liquid and solid media (1% tryptone, 0.5% yeast extract, 1% NaCl, and 1.6% Bacto agar for solid media). Bacterial cultures were grown at 37°C for all experiments or 30°C for strains carrying temperature-sensitive plasmids, unless otherwise specified. Where required, media was supplemented with kanamycin, carbenicillin, chloramphenicol, or tetracycline at a final concentration of 50 μg/ml, 100 μg/ml, 30 μg/ml, or 15 μg/ml, respectively.

### Genetic manipulations

All strains used in study are derivatives of *Salmonella enterica* serovar Typhimurium LT2. The sequences of oligonucleotides (IDT) used in genetic manipulation of strains are provided in **Table S2**. Gene knockouts were created using the λRed method of Datsenko and Wanner using the pKD46 recombinase helper plasmid and site-specific PCR fragments with homologous overhangs generated using pKD13 or pKD32 plasmids as templates (60). Deletion mutants were selected for on LB-kanamycin or LB-chloramphenicol agar plates. All deletion mutants were backcrossed into relevant parental strains using P22 HT/*int* generalized transduction (61). When necessary, antibiotic markers between the FRT sites were excised from mutants by expressing Flp recombinase from pCP20 plasmid (60).

Transcriptional fusions were produced either *in trans* at the λ attachment site or *in situ*. All transcriptional fusions made *in trans* have been described in a previous study and used the pVenus plasmid to produce *yfp* transcription fusions (25, 62). The *in situ* transcriptional fusion to *rflP* used the FLP recombinase-mediated *lacZY* fusion system of Ellermeier and coworkers (63).

### Transposon mutagenesis

Transposon mutagenesis was performed using the T-POP (Tn*10d*Tc[*del-25*]) transposon as described by Lee and coworkers with minor modifications (46, 64, 65). Briefly, high frequency T-POP donor Mu*d*P22 lysates were generated with TH3923 and used to transduce T-POP into LCM2473/pNK2881 cells. Resulting primary T-POP insertion mutants (1,000 - 2,000 colonies/plate) selected on LB-tetracycline agar were pooled, treated with P22 HT *int-105*. In total, 6 independent pools were generated. Random T-POP insertions from each pool were transduced into LCM2473 followed by screening on LB-tetracycline agar plates. Colonies were visually screened for restored *fliC’-yfp* expression using a 400 nm wavelength transilluminator (IORodeo). A minimum of 20,000 T-POP mutants were screened per pool. Positive insertions were mapped by degenerate PCR using primer P-LCM593 to P-LCM598 (**Table S2**) as described by Lee and coworkers (65).

### Transcriptional reporter assays

Transcriptional reporter assays measured either YFP fluorescence or β-galactosidase activity. For YFP fluorescence and β-galactosidase activity assays, cultures were subcultured 1:1000 from overnight cultures and grown to mid-log phase (OD_600_ 0.6-0.8) prior to measurements. To measure YFP fluorescence, 200 μL of each culture was transferred to 96-well black and clear bottom plates (Corning Costar) and fluorescence was measured with a Cytation3 multimode microplate reader (BioTek). Fluorescence was measured at wavelengths 500±5 nm excitation and 525±5 nm emission and normalized to corresponding OD_600_ values. Each sample had a minimum of 6 replicates.

The method of Slauch and Silhavy was used for β-galactosidase activity assays in microtiter plates, with minor modifications (66). Samples (1 mL) from mid-log phase cultures were pelleted at 6,000 x*g* and resuspended in Z buffer (0.1 M sodium phosphate, 0.01 M KCl, 0.001 M MgSO_4_, 0.05 M β-mercaptoethanol, pH 7.0). Two 100 μL aliquots were saved to measure OD_600_ prior to sample permeabilization. Samples were permeabilized via vortexing with 50 μL 0.1% sodium dodecyl sulfate (SDS) and 100 μL chloroform for 1 min. From each sample, two 100 μL aliquots of permeabilized cell extract was added to the wells of a 96 well assay plate. To initiate assays, 20 μL of *o*-nitrophenyl-β-D-galactoside (ONPG) reagent (20 mg/ml in Z buffer) was added followed by mixing at 600 rpm on a microplate shaker (Thermo Scientific). The change in absorbance at 420 nm was measured every 5 min for 1 hr in a Cytation3 multimode plate reader (BioTek). The relative Vmax for each reaction was calculated using Gen5 software (BioTek). β-galactosidase activity of cultures calculated as: Vmax / (OD_600_ * 0.1 mL). Each sample had a minimum of 4 replicates. All data analysis and plotting were performed in R.

### Motility Assays

All motility assays used cultures grown to mid-log phase (OD_600_ 0.6 - 0.8) that were normalized to the lowest measured cell density. For each assay, 1 μL of culture was inoculated into motility agar plates and incubated for 6 hours at 37°C. Images were obtained with a high-resolution flatbed scanner (Epson) and were processed using ImageJ.

### Immunoblots

Immunoblots for FliC, FlhC-3xFLAG, and DnaK were performed with 100 μg of total protein harvested from mid-log phase (OD_600_ 0.6-0.8) cultures per the procedure described elsewhere (25).

### Transcription start site mapping

A template-switching 5’ rapid amplification of cDNA ends (5’-RACE) kit (New England BioLabs) was used to map the transcription start site of *rflP*. Total RNA extracted from mid-log phase cells using TRIzol reagent (Life Technologies) and Direct-zol RNA recovery kits (Zymo Research). cDNA was prepared from total RNA using the *rflP* specific primer (P-LCM621) and template-switching oligonucleotide (P-LCM619) using RT Template Switching Enzyme Mix (New England Biolabs). The 5’-end end of the *rflP* cDNA was enriched by PCR using the primers P-LCM620 and P-LCM622. The enriched fragment is gel purified and sequenced (Eurofins). Sequencing data was mapped to the *S*. Typhimurium LT2 genome using the Artemis software suite.

### Bioinformatic analysis

SoxS binding sites were predicted using the multiple EM for motif elicitation (MEME) algorithm (50). A set of 29 *E. coli* promoters with annotated SoxS binding sites were used to guide the detection of a SoxS binding motif in the *rflP* promoters of *E. coli* and *S*. Typhimurium (**Table S3**). The number of motifs the tool was allowed was set to 1, and the minimum and maximum width of the motif size allowed was set at 12bp and 30bp, respectively. The *rflP* promoter regions of *S*. Typhimurium, and its corresponding region in *E. coli*, were compared using Multiple Sequence Comparison by Log-Expectation tool (MUSCLE) (67).

## ACKNOWLEDGEMENTS

We thank C.M. Cenzer for technical assistance in transposon screening and mapping. This work was supported by NIH grant AI137984 to L.M.C.

